# The GlyGly-CTERM domain functions as an independent motif that targets proteins to rhombosortase in *Vibrio cholerae*

**DOI:** 10.64898/2026.02.05.703148

**Authors:** Cameron S. Roberts, Kwame Kannatey-Asibu, Eliese Potocek, Maria Sandkvist

## Abstract

*Vibrio cholerae* secretes a variety of effector proteins that are freely released into the extracellular space via its Type II Secretion System (T2SS) including cholera toxin, the causative agent of the disease cholera. In contrast to cholera toxin, a growing number of T2SS effectors is increasingly understood to remain associated with the cell surface. The serine protease VesB from *V. cholerae* is a surface protein that is produced with a short C-terminal motif, called GlyGly-CTERM. This motif is linked to the rest of VesB via a predicted unstructured linker. In addition to VesB, *V. cholerae* encodes five additional GlyGly-CTERM proteins including the serine proteases VesA and VesC, a putative metalloprotease VCA0065, the DNase Xds, and VC1485, a protein of unknown function. Proteins with a GlyGly-CTERM are co-distributed in bacteria with a specific rhomboid protease called rhombosortase (RssP), and it has been demonstrated that VesB requires processing by RssP for surface localization and activation. Here, we investigate the intrinsic function of the GlyGly-CTERM by proteomics, enzyme assays and heterologous expression of alternative motifs on model protein VesB as well as on unrelated periplasmic and extracellular proteins. We show that the GlyGly-CTERM and processing by RssP is sufficient for membrane association, but a secondary secretion signal is required for outer membrane translocation. Unexpectedly, VesC is released from the cells through autoproteolytic processing at a site within the unstructured linker. We propose that the GlyGly-CTERM facilitates efficient secretion of proteins via its intrinsic ability to target them to RssP resulting in membrane association.

**Importance:** *Vibrio cholerae* is responsible for the disease cholera. Without treatment, *V. cholerae* causes massive dehydration with high mortality rates (1). It utilizes the Type II Secretion System to export the causative agent of disease, cholera toxin, as well as a suite of additional effector proteins that are involved in pathogenesis. Here, we investigate the unique transport mechanism of a subset of effectors secreted by this pathogen that are targeted to the cell surface.

## Introduction

*Vibrio cholerae* uses the Type II Secretion System (T2SS) to secrete a variety of hydrolytic enzymes into the extracellular space that help the pathogen survive in the environment and colonize the human host (2). Examples of completely secreted effectors from *V. cholerae* include cholera toxin, the driver of the disease cholera, and hemagglutinin/protease (HapA), a metalloprotease capable of degrading the mucosal barrier (3). Increasingly, it is also understood that the T2SS transports proteins to the cell surface (4-9). Multiple mechanisms have been described for cell surface retention of T2SS substrates. For example, pullulanase from *Klebsiella pneumoniae* and InvL of *Acinetobacter baumannii* are modified at their N-terminal lipobox resulting in lipidation prior to outer membrane translocation and cell surface attachment (9, 10). The pectate lyase PnlH is produced with a unique, uncleavable signal peptide that keeps it on the cell surface of *Dickeya dadantii* (5). PdaA has been characterized as a polysaccharide deacetylase that is targeted to the cell surface of *Legionella pneumophila*, but the mechanism of surface association in this case is unknown (7). The serine protease VesB secreted by *V. cholerae* requires the presence and subsequent modification of a C-terminal motif called GlyGly-CTERM for retention on the cell surface (6). This motif is found on a variety of proteins in *V. cholerae* and other gram-negative bacteria such as *A. baumannii* (11). It consists of a glycine rich region, a single pass transmembrane domain (TMD), and a positively charged tail that is connected to the passenger domain via an unstructured linker (6, 11, 12). In the case of VesB, a proteolytic cleavage C-terminal to the double glycine by the rhomboid protease RssP, followed by a possible glycerophosphatidylethanolamine posttranslational modification mediates its surface association following transport across the other membrane by the T2SS (6). *V. cholerae* expresses five additional GlyGly-CTERM proteins, some of which have been shown to rely on the T2SS for secretion (13). In addition to co-distribution of r*ssP* and genes coding for GlyGly-CTERM proteins across genomes, there is also an association with genes for core components of the T2SS indicating a connected activity or function (11). When RssP is absent, VesB is cleaved within its unstructured linker by another rhomboid protease, GlpG, and VesB is no longer targeted to the cell surface. Instead, it is freely released into the extracellular space after outer membrane translocation (12).

GlyGly-CTERM proteins expressed by *V. cholerae* include VesB and two related serine proteases, VesA and VesC, a putative metalloprotease, VCA0065, the DNase Xds, and a protein of unknown function, VC1485. VesA, VesB and VesC have all been characterized as secreted substrates of the T2SS, but additional GlyGly-CTERM proteins have yet to be confirmed as dependent on the T2SS for secretion (13). Trypsin-like specificity has been described for VesB and VesC (13, 14). Based on the sequence of its predicted propeptide, VesA is likely a chymotrypsin-like serine protease (13). Both VesA and VesB have been shown to cleave the A-subunit of cholera toxin, a process required for activation of the toxin (13). VesC induces a hemorrhagic fluid response when injected into a mouse ileal loop system (15). Studies on Xds suggest that it supports evasion of neutrophil extracellular traps, presumably through its DNase activity (16). While surface localization of VesB has been demonstrated (6, 9, 12), it is unclear if the other GlyGly-CTERM-containing proteins are also targeted to the cell surface. One of the GlyGly-CTERM proteins expressed by *V. cholerae*, VesC, is toxic to the cell in T2SS mutants (17). Construction of T2SS deletion mutants often results in the selection of inactivating mutations in VesC, and overexpression of active VesC in these mutant strains is lethal (17). Despite their homology with VesC, inactivating suppressor mutations have not been identified in VesA or VesB of T2SS mutants. While the genomes of an exhaustive list of T2SS mutants have yet to be sequenced, inactivating mutations have not been detected in any other known T2SS effector either, perhaps indicating a unique mechanism of VesC toxicity when it accumulates in the periplasmic space of T2SS mutants.

To further elucidate the role of the GlyGly-CTERM and determine whether it functions independently of its passenger domain, we engineered chimeric VesB constructs with the GlyGly-CTERM from VesA, VesC, and VC1485 and determined the impact on VesB secretion and activation. We also genetically fused the GlyGly-CTERM from VesB to the C-terminus of other proteins that are normally destined for the periplasmic space or culture supernatant and determined their subcellular location. In addition, we probed the cellular location of all GlyGly-CTERM proteins in *V. cholerae* using an unbiased quantitative proteomic approach. Our results indicate that the GlyGly-CTERM is a transferable RssP targeting motif that harbors ‘plug-and-play’ capability. Expression of proteins with this motif and a separate secretion signal recognized by the T2SS support their surface localization.

## Results

### The GlyGly-CTERM from VesA, VesC, and VC1485 are interchangeable with that of VesB

We have previously demonstrated that VesB depends on RssP for activation and that VesB expressing an alternative transmembrane domain (TMD) is not recognized by RssP and thus not activated (12). While the GlyGly-CTERM motifs found in *V. cholerae* share some similarity, there are appreciable differences in their amino acid composition (**Figure 1A**). To determine the effect of expressing VesB with alternative GlyGly-CTERM motifs, we engineered constructs replacing residues C-terminal to the di-glycine motif of VesB (G382-G383) with those from VesA, VesB and VC1485 (**Figure 1B**). Expression of VesB with the alternative GlyGly-CTERM motifs was sufficient to generate active, cell associated VesB, indicating proper recognition by RssP (**Figure 1C**). Indeed, when we modeled the interaction of the VesB chimeric proteins with RssP they adopted a similar confirmation to that of native VesB where the TMD of each protein was docked into the canonical rhomboid protease substrate recognition site between TMD2 and 5 (**Figure 1D, Sup. Figure 1**). Notably, no activity was detected in the culture supernatant for VesB expressed with the GlyGly-CTERM of VesC, while the other chimeras showed high extracellular activity similar to that of native VesB.

**Figure 1.**
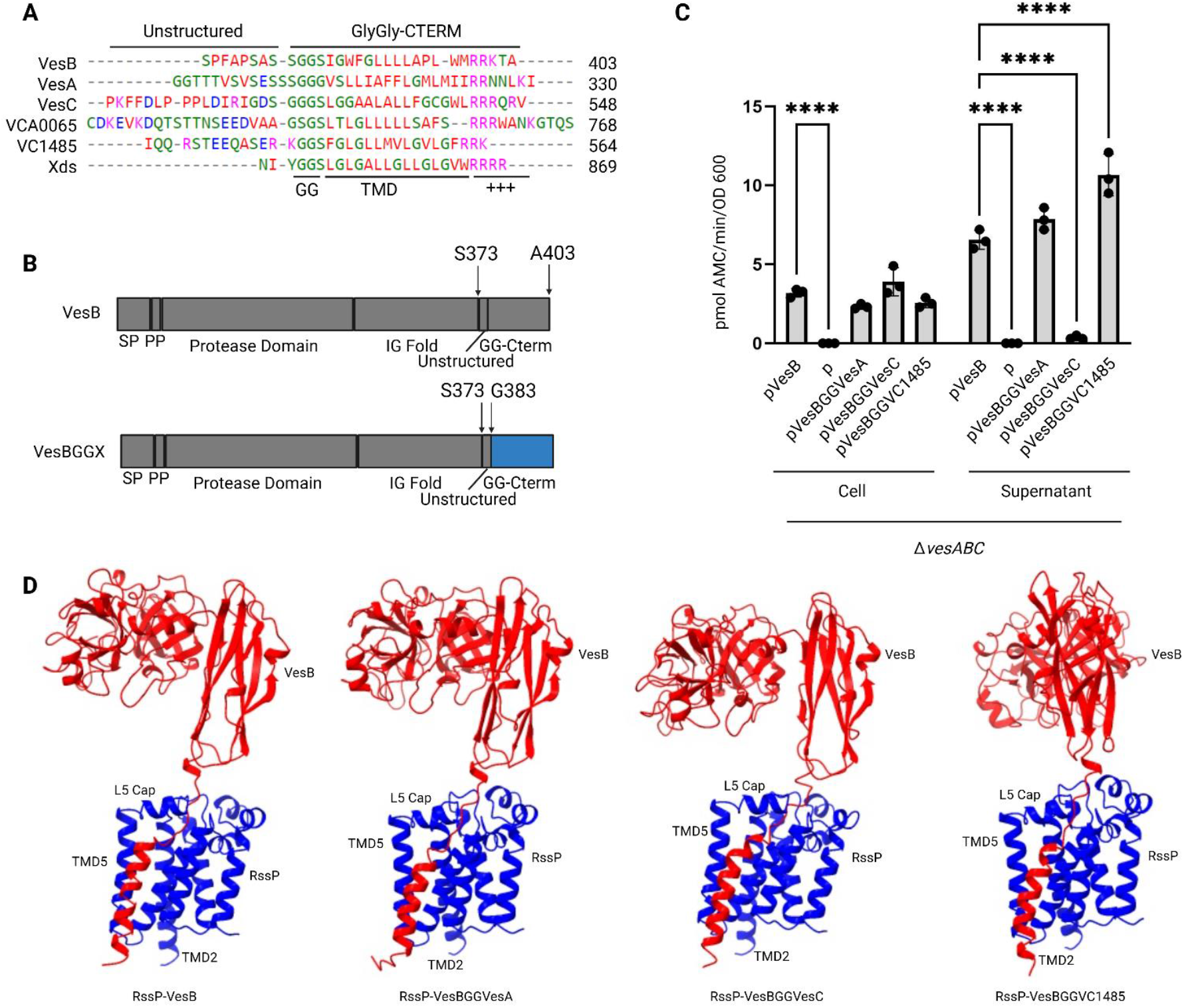
GlyGly-CTERM motifs are functionally interchangeable. **A.** Alignment of amino acid sequence of the GlyGly-CTERM and the preceding unstructured region of six *V. cholerae* proteins including VesA, VesB, and VesC. Unstructured region, as predicted by AlphaFold, glycine signal (GG), transmembrane domain (TMD), as predicted by DeepTMHMM, and positively charged tail (+++) are schematically indicated. **B**. Schematic organization of VesB and VesB chimeric proteins containing the GlyGly-CTERM (indicated in blue) from VesA, VesC, or VC1485 with Signal peptide (SP), Propeptide (PP), and GlyGly-CTERM (GG-CTERM) specified. **C**. Overnight cultures of the mutant strain Δ*vesABC* containing empty vector (p) or plasmids encoding WT VesB or indicated VesB chimera were separated into cell and culture supernatant fractions and assessed for serine protease activity using the fluorogenic peptide substrate Boc-Gln-Ala-Arg-AMC. Data represent mean ± SD of *n* = 3 experiments done in technical triplicate. ****P≤0.0001; two-way Anova multiple comparison test with Sidak post hoc analysis. Only significant differences are indicated. **D**. A predicted interaction of RssP (blue) and VesB or VesB chimera (red) was generated using AlphaFold3. Transmembrane domain (TMD) and loop 5 (L5) cap of RssP indicated. Note that in the predicted interactions we have oriented the structure relative to RssP resulting in a slight rotation of VesBGGVC1485 with respect to RssP and the other predictions.

### Proteomic analysis reveals differential localization of GlyGly-CTERM proteins

Given our new and previous findings (6, 12) with VesB and VesB chimeric proteins, we wanted to investigate whether the other five GlyGly-CTERM containing proteins in *V. cholerae* also localize to the membrane fraction, and likely cell surface, in a RssP-dependent manner. If the GlyGly-CTERM motifs of these proteins function similarly to that of VesB, we expected to detect these proteins in the membrane fraction of WT cells, but not of *rssP*::kan mutant cells by proteomics. When we compared the membrane protein profiles of WT and *rssP*::kan mutant cells, we detected peptides corresponding to VesB and Xds exclusively in the WT preparations, and an enrichment of peptides for VesA and VC1485 that was statistically significant (**Figure 2A, Table 1**). As a control, we compared the profiles of WT and Δ*glpG* mutant membrane fractions and did not observe any statistically significant differences for the GlyGly-CTERM proteins (**Figure 2B**). We did observe higher peptide counts for VesB in the WT membrane fraction as compared to the Δ*glpG* mutant, but this difference was not statistically significant. These findings validated that VesB, and likely Xds, VesA, and VC1485, are localized to the membrane fraction in an RssP dependent manner. We did not detect peptides for VCA0065 or VesC across any membrane preparation. To determine if either protein was present in the culture supernatant we also subjected culture supernatants from WT and rhomboid mutant cells to proteomic analysis (**Figure 2C** and **D**). This revealed that the amount of VesC was significantly higher in the culture supernatant of the WT strain than that of the *rssP*::kan mutant. VesA, Xds, and VCA0065 were also more abundant in WT supernatants as compared to the supernatant of the *rssP*::kan strain, although lower peptide counts meant this result did not reach statistical significance. VesB and VC1485 were trending to be enriched in *rssP*::kan mutant supernatants as compared to the WT strain, indicating that these proteins are released to the extracellular space in the absence of RssP. While it has yet to be confirmed for VC1485, we previously demonstrated that processing of VesB by GlpG results in the release of VesB into the culture supernatant in the absence of RssP (12). Indeed, when we compared the amount of these proteins released to the culture supernatant from the *rssP*::kan mutant strain to the Δ*glpG* mutant strain, we observed an enrichment in the *rssP*::kan supernatants of VesB and VC1485 (**Table 1**).

**Table 1.** Label free proteomics identifies GlyGly-CTERM proteins in WT and rhomboid mutant strains.

| Supernatant |  |  |  |  |  |  |
| --- | --- | --- | --- | --- | --- | --- |
| | WT<br>Identified<br>Peptides | *rssP::kan*<br>Identified<br>Peptides | Δ*glpG*<br>Identified<br>Peptides | WT NSAF<br>/*rssP::kan*<br>NSAF | WT<br>NSAF/Δ*glpG*<br>NSAF | *rssP::kan*<br>NSAF/Δ*glpG*<br>NSAF |
| **VCA0065** | 42±13 | 33±17 | 38±14 | 2.8 | 1.1 | 0.4 |
| **VesC** | 60±13 | 31±11 | 57±14 | 7.3 | 0.7 | 0.1 |
| **Xds** | 55±15 | 35±24 | 51±12 | 3.7 | 1 | 0.2 |
| **VesA** | 10±1 | 7±5 | 11±0 | 4.8 | 1.2 | 0.2 |
| **VesB** | 45±12 | 63±12 | 30±6 | 0.7 | 1.5 | 2.1 |
| **VC1485** | 54±10 | 52±10 | 69±5 | 0.4 | 0.8 | 2.2 |
| Membrane |  |  |  |  |  |  |
| | WT<br>Identified<br>Peptides | *rssP::kan*<br>Identified<br>Peptides | Δ*glpG*<br>Identified<br>Peptides | WT NSAF<br>/*rssP::kan*<br>NSAF | WT<br>NSAF/Δ*glpG*<br>NSAF | *rssP::kan*<br>NSAF/Δ*glpG*<br>NSAF |
| **VCA0065** | 0±0 | 0±0 | 0±0 | n/a | n/a | n/a |
| **VesC** | 0±0 | 0±0 | 0±0 | n/a | n/a | n/a |
| **Xds** | 8±6 | 0±0 | 6±4 | 5* | 1.1 | 0 |
| **VesA** | 22±8 | 1±1 | 19±9 | 380.8 | 1.2 | 0 |
| **VesB** | 1±0 | 0±0 | 2±2 | 5* | 0.4 | 0 |
| **VC1485** | 104±13 | 7±5 | 108±16 | 12.5 | 1 | 0.1 |

Average of three experiments shown for supernatant and membrane fractions.
\*Indicates arbitrary fold change assigned for proteins identified in WT samples but not *rssP::kan*.

**Figure 2.**
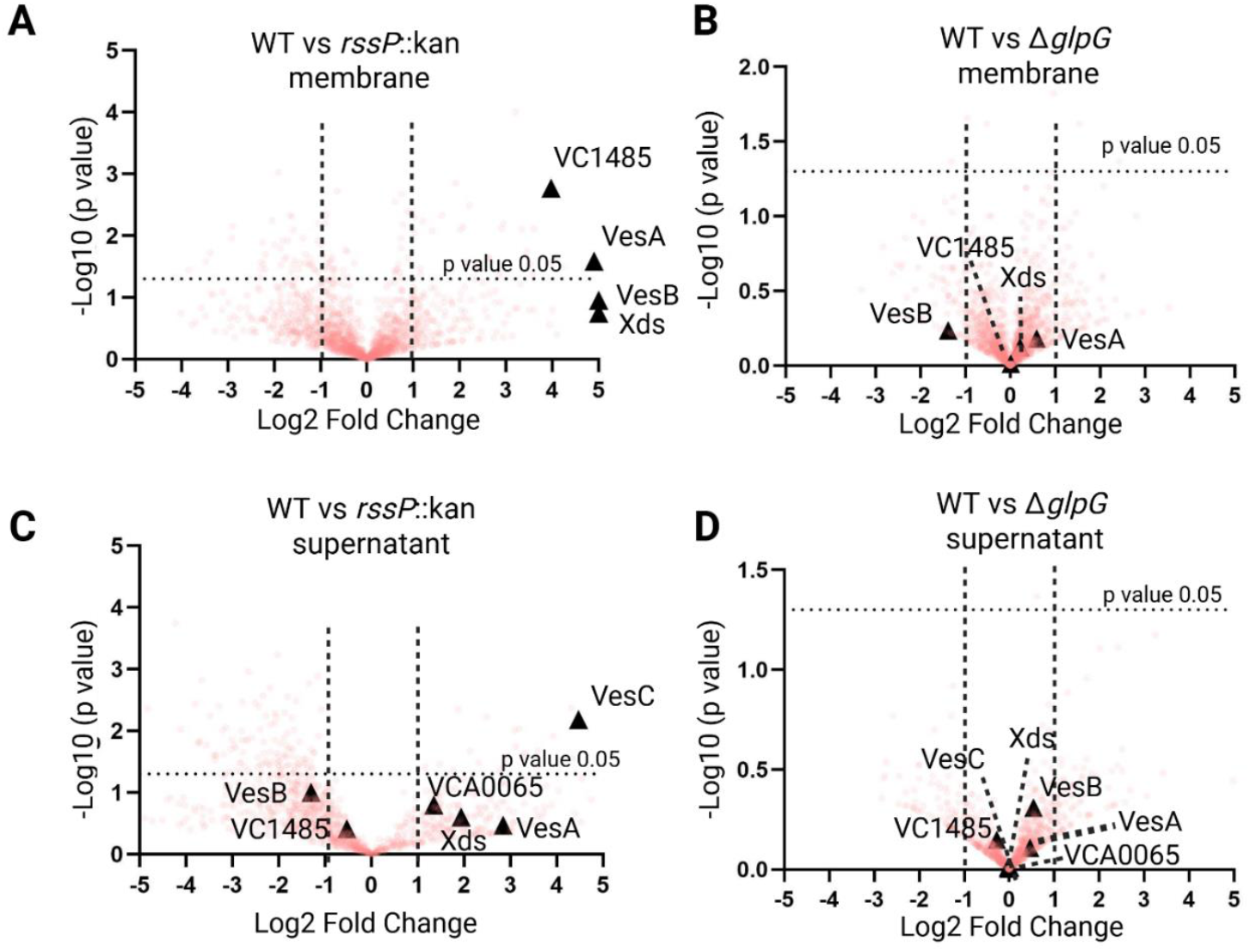
GlyGly-CTERM proteins are differentially localized across the membrane and culture supernatant in an RssP-dependent manner. **A**. and **B**. Volcano plots depicting LC/MS-MS data of proteins identified in isolated membranes of WT versus *rssP*::kan mutant (**A**) or WT versus Δ*glpG* mutant (**B**) strains (n = 3). **C**. and **D**. Volcano plots depicting LC/MS-MS data of proteins identified in isolated culture supernatants of WT versus *rssP*::kan mutant (**C**) or WT versus Δ*glpG* mutant (**D**) strains (n = 3). Log2-fold change of the Normalized Spectral Abundance Factor (NSAF) is shown on the x-axis against statistically significance (p value) determined by student t-test on the y-axis. A value of -5 was manually assigned to proteins only detected in mutant strains, where a value of 5 was assigned to proteins only detected in WT strain. GlyGly-CTERM proteins are indicated.

As noted above, we observed that VesC was highly enriched in the WT supernatant samples as compared to the *rssP*::kan mutant cells despite not detecting any VesC peptides in the membrane fraction across any strain. To validate if active VesC is preferentially secreted in a RssP dependent fashion, we utilized the ActivX serine hydrolase probe, which labels serine hydrolases including VesA, VesB and VesC once they are activated (**Sup. Figure 2A**). After fractionation of WT and Δ*vesC* mutant cells into culture supernatant, periplasmic, cytoplasmic, and membrane fractions we determined that native, active VesC was primarily found in the culture supernatant (**Sup. Figure 2B**). We then subjected WT and rhomboid mutant supernatants to ActivX labeling and observed that active VesC was secreted in an RssP-dependent manner (**Sup. Figure 2C**). We subsequently validated these findings by ectopically overexpressing VesC in WT and rhomboid mutant backgrounds (**Sup. Figure 2D** and **E**). While processing by RssP is required for subsequent activation of VesB under native conditions, we were previously able to detect activity of VesB when overexpressed in a *rssP* mutant strain (12). In contrast, secreted active VesC was not detected when *rssP* was inactivated even under overexpression conditions. Given our findings with these two serine proteases, we turned to the homologous protease VesA and asked whether its presence as an active protease in the culture supernatant is dependent on one or both rhomboid proteases. Using the fluorogenic peptide Suc-Leu-Leu-Val-Tyr-AMC, which is primarily a substrate for VesA in WT *V. cholerae* (**Sup. Figure 3A**), we assessed native VesA activity in the culture supernatants of WT and rhomboid mutant cells. Like both VesB and VesC, active VesA was only detected in culture supernatants in the presence of active RssP (**Sup. Figure 3B**). Furthermore, similar to VesC, ectopic overexpression of VesA resulted in activity detected in WT and Δ*glpG* mutant cells, but not in *rssP*::kan mutant cells (**Sup. Figure 3C**).

**Figure 3.**
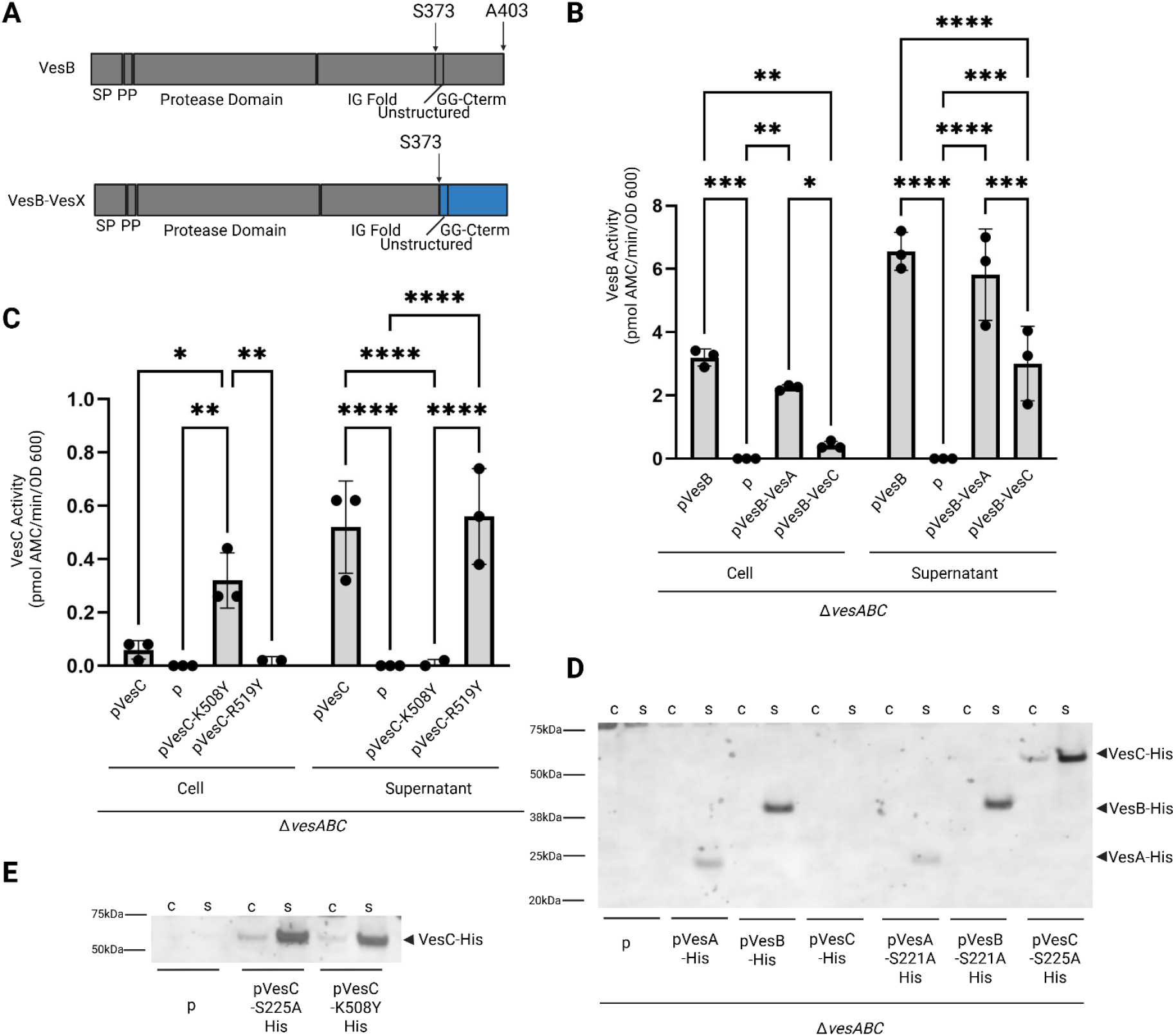
The disordered region of VesC is targeted for proteolysis. **A.** Schematic organization of VesB (top) and VesB chimeric proteins containing the GlyGly-CTERM and unstructured region (indicated in blue) from either VesA or VesC (bottom). Signal peptide (SP), propeptide (PP), and GlyGly-CTERM (GG-CTERM) are shown. **B**. Overnight cultures of the mutant strain Δ*vesABC* containing empty vector (p) or plasmids encoding WT VesB or indicated VesB chimera were separated into cell and culture supernatant fractions and assessed for serine protease activity using the fluorogenic peptide substrate Boc-Gln-Ala-Arg-AMC. Data represent mean ± SD of *n* = 3 experiments done in technical triplicate. * representing p<0.05, **p<0.005, ***p<0.0005, and **** p≤0.0001; two-way Anova multiple comparison test with Sidak post hoc analysis. Only statistically significant differences are indicated. **C**. Overnight cultures of the mutant strain Δ*vesABC* containing empty vector (p) or plasmids encoding WT VesC or indicated VesC mutant protein were separated into cell and culture supernatant fractions and assessed for serine protease activity using the fluorogenic peptide substrate Boc-Gln-Ala-Arg-AMC. Data represent mean ± SD of *n* = 3 experiments done in technical triplicate. * representing p<0.05, **p<0.005, ****p≤0.0001; two-way Anova multiple comparison test with Sidak post hoc analysis. Only statistically significant differences are indicated. **D**. Overnight cultures of the mutant strain Δ*vesABC* containing empty vector (p) or plasmids overexpressing (10 µM IPTG) VesA, VesB, or VesC mutant proteins were separated into cell (c) and culture supernatant (s) fractions, run on SDS-PAGE, transferred to nitrocellulose membrane, and blotted with anti-His6 antibodies. VesA, VesB, and VesC species are indicated. Representative blot is shown (n=2). **E**. Overnight cultures of the mutant strain Δ*vesABC* containing empty vector (p) or plasmids overexpressing (10 µM IPTG). VesC mutant proteins were separated into cell (c) and culture supernatant (s) fractions, run on SDS-PAGE, transferred to nitrocellulose membrane, and blotted with anti-His6 antibodies. VesC species are indicated. Representative blot is shown (n=2).

Given the lack of VesC detected in the membrane fraction of WT cells, we were curious if any amount of VesC released to the culture supernatant was associated with the pelletable fraction containing OMVs. When the culture supernatants from Δ*vesABC* mutant cells ectopically overexpressing VesB or VesC were subjected to high-speed centrifugation, we observed a higher percentage of VesB activity associated with the pelletable fraction compared to VesC activity suggesting that VesB can be released in association with OMVs whereas VesC is primarily released from the cells in a soluble form (**Sup. Figure 2F**). Collectively, these results indicate that unlike VesB, VesC is primarily secreted, is not processed by GlpG, and is fully released into the culture supernatant.

### VesC is transiently retained by the cell before proteolytic release

To explain why VesC is primarily detected in the culture supernatant despite finding the VesBGGVesC chimera mostly associated with the cell pellet, we examined the predicted unstructured region upstream of the GlyGly-CTERM. Previously we observed that this linker region of VesB is subject to proteolytic cleavage by GlpG in the absence of RssP, and substitutions of residues with those present in the unstructured region of VesC blocked GlpG cleavage (12), explaining the lack of GlpG processing of VesC observed in this study. We hypothesized that this same region in VesC might be subject to cleavage by another protease resulting in the release of VesC to the culture supernatant. We therefore generated an additional chimeric VesB protein containing both the linker region and GlyGly-CTERM from VesC as well as the equivalent residues from VesA as a control (**Figure 3A**). When VesB was expressed with its native sequence or that from VesA, we detected activity in both cell and supernatant fractions using the fluorogenic peptide substrate Boc-Gln-Ala-Arg-AMC (**Figure 3B**). In contrast, when we expressed VesB with the VesC sequence, VesB activity was largely detected in the culture supernatant demonstrating that the unstructured region and GlyGly-CTERM from VesC is sufficient to support the full release of VesB to the culture supernatant. Because VesB-VesC, and VesBGGVesC, both appear to have lower total activity as compared to WT VesB, we compared the ratio of cell to supernatant activity of cultures expressing each construct using the VesA mutants as a control (**Table 2**). We observed a greater than 30-fold increase in the relative cell associated activity of VesBGGVesC as compared to VesB, while the VesB-VesC mutant displayed a greater than 3-fold decrease in cell associated activity. To determine if the VesB-VesC chimera is transiently associated with cells, we performed a time course experiment monitoring the cell and supernatant VesB activity (**Sup. Figure 4**). We observed that initial time points indicate that VesB-VesC is transiently associated with the cells similarly to VesB and VesB-VesA, and likely the cell surface given our previous findings on VesB activation, but a greater fraction is subsequently released from the cells (6, 9). We reasoned that VesC, or the VesB-VesC chimera, may cleave itself once targeted to the cell surface and given its trypsin-like specificity that this might occur at lysine 508 or arginine 519 in the predicted unstructured region that precedes GlyGly-CTERM. To determine the effect on localization of VesC we separately substituted K508 and R519 with tyrosine and assessed cells and supernatants fractions for protease activity using the fluorogenic substrate (**Figure 3C**). Activity of both WT VesC and VesC-R519Y was primarily detected in the culture supernatant, while VesC-K508Y activity was found in the cell fraction, suggesting that this mutation blocked proteolytic cleavage and release from the cells reflecting our results with the VesBGGVesC chimera that also lacks this proteolytic site. Analysis of the ratio of activity in cell and supernatant fractions again demonstrate that these observations were not driven by an overall reduction in activity (**Table 2**). To further determine if VesC undergoes proteolytic cleavage at its C-terminus, we took advantage of previously engineered mutant constructs of VesA, VesB, and VesC, each with the catalytic serine substituted with an alanine and the residues C-terminal of the double glycine in the GlyGly-CTERM motif replaced with a His tag (9). These constructs were completely released to the culture supernatant (9). Here we generated a second set of His-tagged constructs, but using the three proteases with their catalytic serine intact. When ectopically expressed, we detected VesA-His and VesB-His in the culture supernatants of *vesABC* mutant cells by western blotting using anti-His antibodies regardless of whether their catalytic serine residue was substituted with alanine or not (**Figure 3D**). In contrast, we only detected the His tag of the catalytically inactive VesC-His mutant, suggesting that VesC undergoes auto-proteolysis at its C-terminus. Indeed, we detected activity associated with VesC-His in the culture supernatant (**Sup. Figure 5**). We also subjected the VesC-His construct to mutagenesis at position K508, to determine if the K508Y mutation again blocked C-terminal processing. When we expressed active VesC-K508Y His or VesC-S225A His we detected a band corresponding to VesC in the culture supernatant via western blotting for each construct (**Figure 3E**). The lack of C-terminal processing by VesC-K508Y-His could not be explained by a lack of activity as it demonstrated similar levels of activity compared to VesC-His using the fluorogenic peptide substrate Boc-Gln-Ala-Arg-AMC (**Sup. Figure 5**). To assess if the C-terminus of VesC was processed in trans by VesA, VesB, or VesC, we expressed inactive VesC-His in the presence of native VesA, VesB, or VesC (**Sup. Figure 6**). We did not observe appreciable loss of His detection regardless of protease background.

**Table 2.** Relative cell to supernatant activity of VesB, VesC, indicated chimeras, and VesC mutant proteins.

| Protein | Ratio Cell/Supernatant<br>Activity |
| --- | --- |
| VesB | 0.49 ± 0.07 |
| VesBGGVesA | 0.30 ± 0.03 |
| VesBGGVesC | 16.33 ± 3.76 |
| VesB-VesA | 0.41 ± 0.09 |
| VesB-VesC | 0.15 ± 0.04 |
| VesC | 0.13 ± 0.08 |
| VesC-K508Y | 12.33 ± 0.94 |
| VesC-R519Y | 0.05 ± 0.02 |

**Figure 4.**
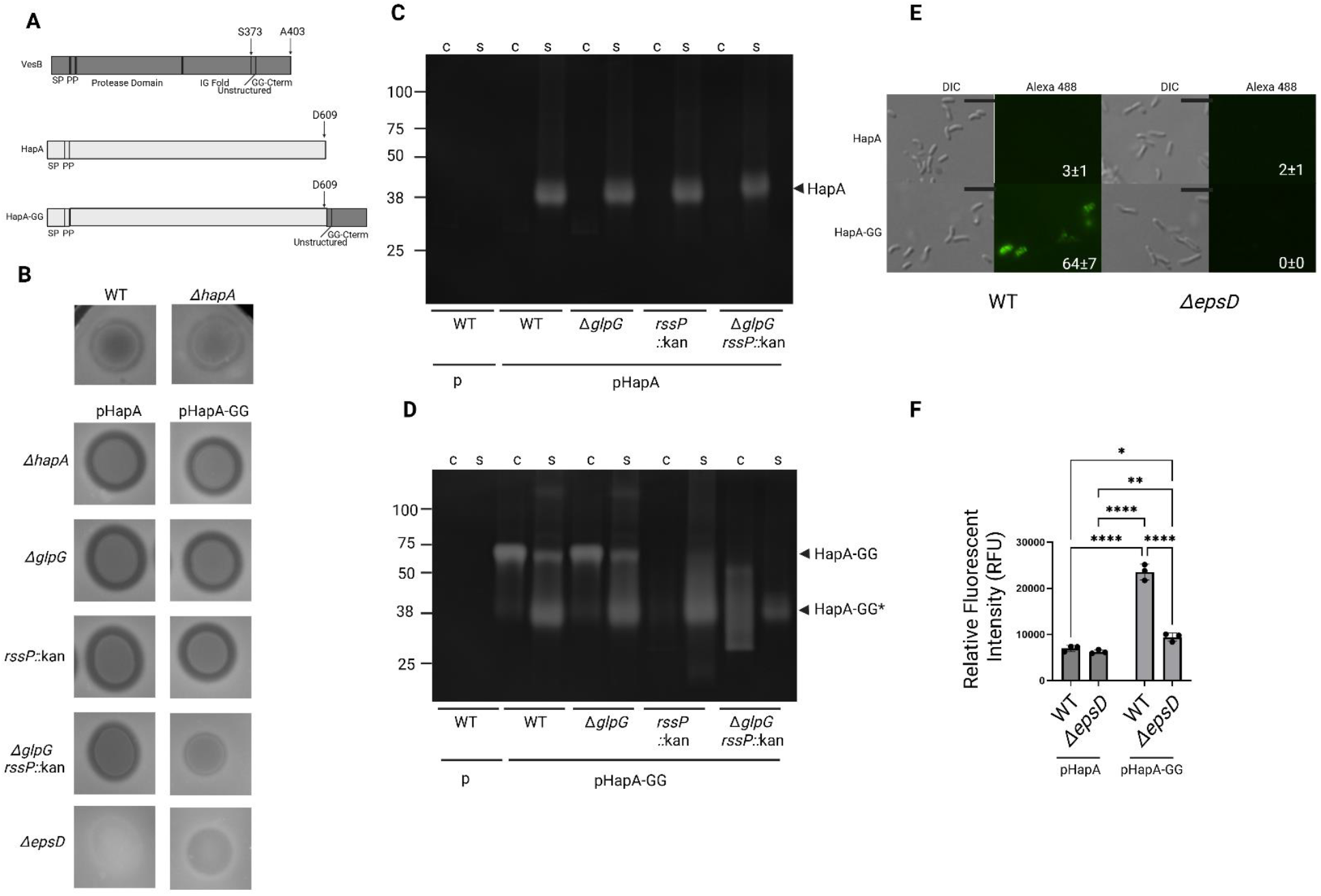
HapA is processed by the rhomboid proteases RssP and GlpG and localizes to the cell surface when fused with the GlyGly-CTERM from VesB. **A**. Schematic organization of VesB, HA/protease (HapA), and chimeric HapA containing the GlyGly-CTERM and unstructured linker (indicated in dark gray) from VesB (HapA-GG). Signal peptide (SP), propeptide (PP), protease domain, immunoglobulin (IG) fold and GlyGly-CTERM (GG-CTERM) are indicated. **B**. (top) Overnight cultures of WT and *hapA* protease mutant strain (Δ*hapA*) were spotted on 1% milk agar plates before incubation overnight at 37 °C and visualization for clearance zones. (bottom) Overnight cultures from indicated mutant strains ectopically expressing WT or chimeric HapA were spotted on 1% milk agar plates before incubation overnight at 37 °C and visualization of protease activity as indicated by casein clearance. Three independent experiments were performed with representative data shown. **C**. HapA- and **D**. HapA-GG-positive cultures from **B** were separated into cell and supernatant fractions and subjected to SDS-PAGE on zymogram gels containing gelatin before visualizing protease activity in the form of gelatin hydrolysis after removal of SDS and staining with Coomassie brilliant blue. Representative gel is shown (n=3). Arrows indicate the size expected of mature HapA and a higher molecular weight species, HapA-GG. **E**. Cells from log phase cultures of WT or Δ*epsD* mutant cells ectopically expressing HapA or HapA-GG were isolated, incubated with anti-HapA antibodies followed by Alexa 488 conjugated secondary antibody and visualized by fluorescence microscopy (top) and differential interference contrast microscopy (bottom). Representative image is shown from three independent experiments. Bar represents 5 µm. Numbers indicate percentage fluorescent cells (n=3, 100 cells each). **F**. Suspended cells from **E** were measured for fluorescence intensity using excitation/emission wavelengths of 488/519 nm in wells of black microtiter plates. Data represent mean ± standard deviation (SD) of *n* = 3 experiments in technical triplicate. *P<0.05, **p<0.005 and ****p≤0.0001 by two-way ANOVA analysis with a Tukey correction for multiple comparisons. Only significant differences are indicated.

**Figure 5.**
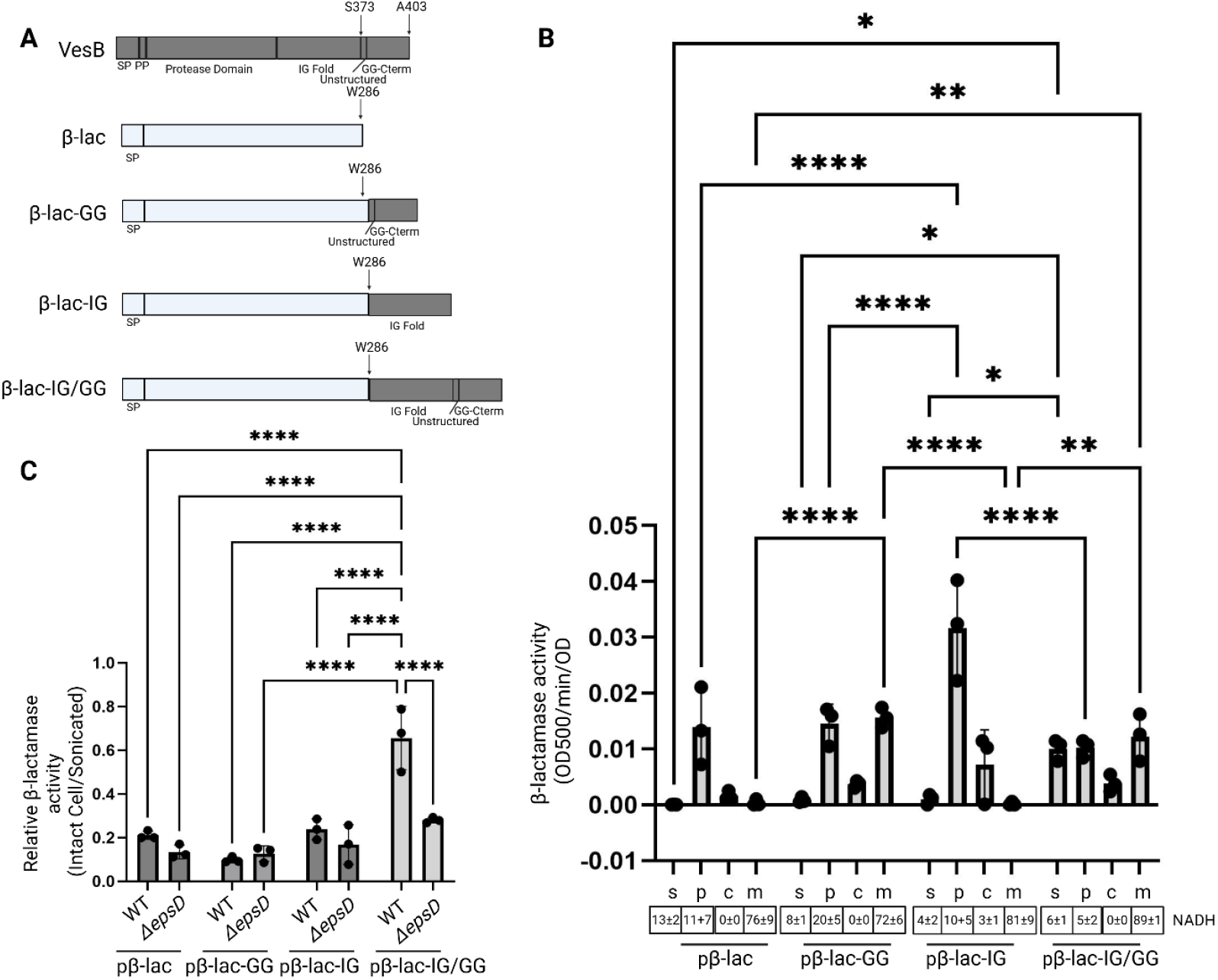
The GlyGly-CTERM works in tandem with a secretion signal to facilitate surface localization of β-lactamase. **A**. Schematic organization of VesB, β-lactamase (β-lac) and indicated β-lactamase chimeras containing the GlyGly-CTERM (GG) with the unstructured linker, immunoglobulin (IG) fold, or both from VesB (indicated in dark gray). Signal peptide (SP), propeptide (PP), and protease domain are indicated. **B**. Log phase cultures of WT cells expressing β-lac or indicated chimera were fractionated into culture supernatant (s), periplasmic (p), cytoplasmic (c), and total membrane (m) fractions, before each fraction was assessed for β-lactamase activity using nitrocefin as a substrate (top). Data represent mean ± SD of *n* = 3 experiments that were analyzed in technical triplicate. *P<0.05, **p<0.005, and ****p≤0.0001 by two-way Anova analysis with a Tukey correction for multiple comparisons. Only significant differences are indicated. As a control for membrane isolation, NADH dehydrogenase activity was monitored by measuring the decrease in NADH absorbance at 340 nm over time. Data represent average ± SD from technical duplicates are shown (bottom). **C**. Cells from log phase cultures of WT or Δ*epsD* mutant cells expressing β-lac or indicated chimera were isolated and intact cells (IC) or sonicated cells were assessed for β-lactamase activity using nitrocefin as a substrate where values are expressed as the ratio of activity of intact cells over sonicated cells. Data represent mean ± SD of *n* = 3 experiments analyzed in technical triplicate. ****p≤0.0001 by two-way Anova analysis with a Tukey correction for multiple comparisons. Only statistically significant differences are indicated.

**Figure 6.**
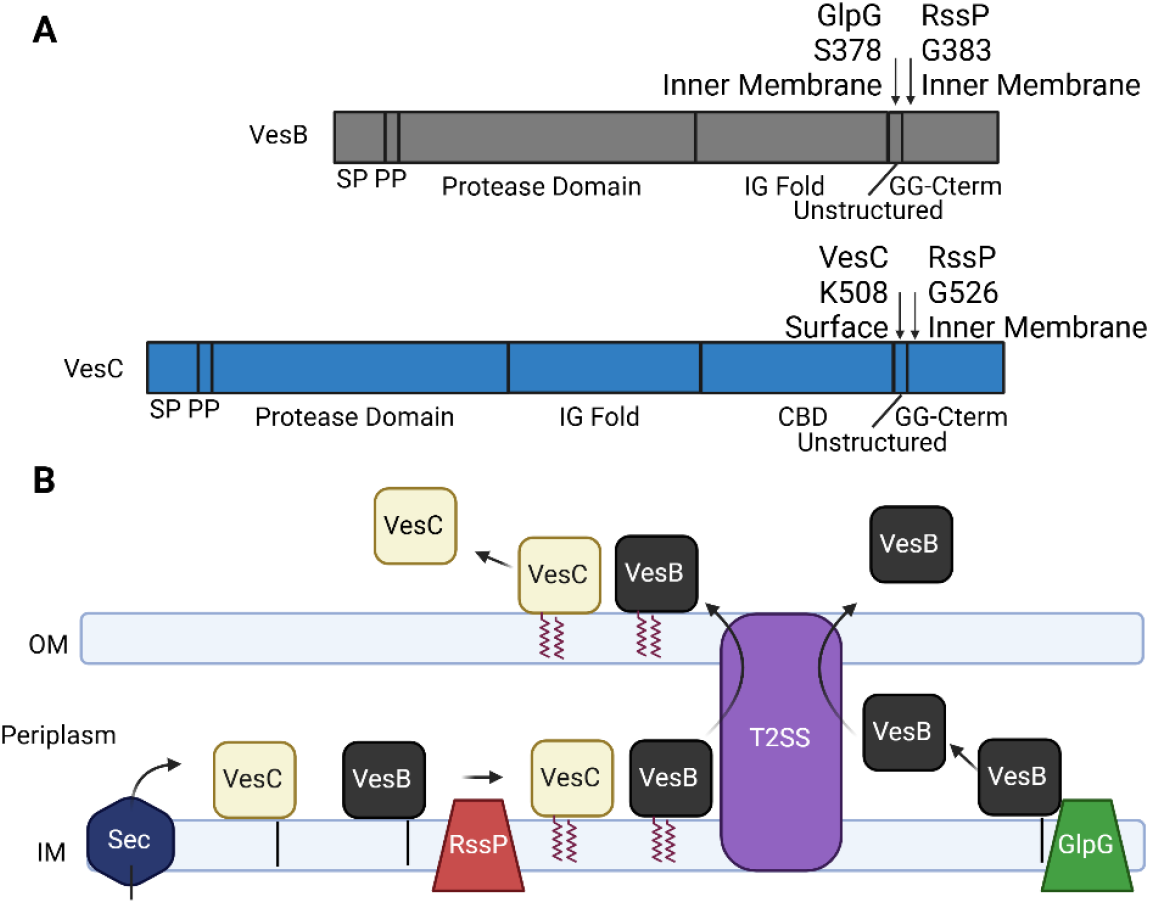
Proposed biogenesis of VesB and VesC. **A.** Schematic organization of VesB and VesC. Signal peptide (SP), propeptide (PP), GlyGly-CTERM (GG-CTERM), IG Fold, and carbohydrate binding domain (CBD) are shown. Arrows indicate proposed proteolytic sites and proposed cellular location of the proteolytic event. **B**. Proposed model of VesB and VesC secretion. Proteins are initially transported into the periplasm by the SEC pathway where VesB and VesC are recognized by RssP and posttranslationally modified before transport to the cell surface via the T2SS. Alternatively in the absence of RssP, VesB is processed by GlpG and released to the culture supernatant after outer membrane translocation. VesC is released from the cell surface through an autoproteolytic event.

We previously demonstrated that VesB requires an intact GlyGly-CTERM and processing by RssP for subsequent removal of its propeptide and activation. To assess the impact of expressing VesC without the GlyGly-CTERM on its activity, we expressed VesC-His under non-inducing (0 µM IPTG) and inducing conditions (10 µM) and observed that overexpression conditions were required for generating similar levels of activity as full length VesC under non-inducing conditions (**Sup. Figure 5;** compared with **Figure 3C**). We also discovered that VesA needs to be expressed with its GlyGly-CTERM in order to be efficiently activated (**Sup. Figure 3D**).

### HapA expressed with the GlyGly-CTERM from VesB is localized to the cell surface

Next, we investigated the possible self-contained function of the GlyGly-CTERM motif addressing if it can heterologously target another T2SS substrate to RssP for processing and subsequent cell surface localization. To this end, we genetically engineered a HapA chimera (HapA-GG) with the C-terminal GlyGly-CTERM and the preceding disordered region from VesB (**Figure 4A**). To monitor the secretion of HapA-GG we spotted cells on agar containing 1% milk and visualized protease activity in the form of clearance of casein after overnight incubation where we utilized *V. cholerae* expressing wild type (WT) HapA as positive control. We used the El tor strain N16961, which has a point mutation in the *hapA* regulator, HapR (18), and therefore has negligible levels of native HapA activity as evident when compared to an Δ*hapA* mutant strain (**Figure 4B**). When WT HapA was ectopically expressed in N16961, we could detect zones of clearance for WT cells, Δ*glpG, rssP*::kan, and Δ*glpG rssP*::kan mutant cells. Consistent with a role for the T2SS in the secretion of HapA, we did not see casein clearance when HapA was ectopically expressed in the T2SS mutant strain Δ*epsD*. Like WT HapA, HapA-GG produced a zone of clearance in WT, Δ*glpG* and *rssP*::kan mutant backgrounds, but not in the Δ*epsD* mutant. In contrast, when HapA-GG was expressed in Δ*glpG rssP*::kan mutant cells, we did not observe casein digestion, which is consistent with previous results demonstrating that VesB requires RssP or GlpG for secretion (12). To distinguish between a fully secreted and a potentially surface localized HapA species, we subjected cell and culture supernatant fractions from WT or rhomboid mutant strains expressing either WT HapA (**Figure 4C**) or HapA-GG (**Figure 4D)** to SDS-PAGE and gelatin zymography as a readout for HapA enzyme activity. We observed that WT HapA was secreted regardless of strain background and traveled at an expected molecular weight of the mature protease species (19). HapA normally undergoes several proteolytic events towards a fully active protease including cleavage of its signal peptide, prodomain, and an unidentified C-terminal event resulting in a mature protease with a molecular weight of approximately 35 kDa (19). In contrast to WT HapA, HapA-GG was partially retained in the cell fraction in an RssP-dependent manner and traveled as a significantly higher molecular weight species (approximately 68 kDa). This higher molecular weight species is consistent with the expected molecular weight of a HapA that lacks processing of its C-terminus. In the Δ*glpG rssP*::kan double mutant cell and culture supernatant fractions the banding pattern was suggestive of HapA-GG degradation similarly to what we have previously observed for VesB when expressed in this rhomboid null mutant (12). To determine if cell-associated HapA-GG is localized on the cell surface similar to VesB, we subjected intact cells to fluorescence microscopy using HapA antibodies and Alexa Fluor 488 conjugated secondary antibodies. We were able to efficiently detect HapA-GG expressed in WT cells, but not in the Δ*epsD* mutant strain (**Figure 4E** and **F)**. Not surprisingly, regardless of background, no cell surface signal was detected for WT HapA. Our results indicate that the GlyGly-CTERM from VesB is capable of successfully targeting HapA to RssP for processing and cell surface localization, demonstrating a plug-and-play functionality.

### The GlyGly-CTERM from VesB works in tandem with a secretion signal to direct a periplasmic enzyme to the cell surface

We previously showed that the Immunoglobulin (IG) fold and GlyGly-CTERM from VesB (**Figure 5A**) is sufficient to guide β-lactamase to the culture supernatant as assessed by monitoring β-lactamase activity using nitrocefin as substrate (9). To dissect the individual contribution of the IG fold and GlyGly-CTERM towards β-lactamase secretion, we engineered two additional constructs consisting of β-lactamase fused to either the IG fold or the GlyGly-CTERM including the unstructured linker region from VesB. We subjected WT *V. cholerae* expressing these constructs to subcellular fractionation and measured the activity of β-lactamase (**Figure 5B**). As compared to normally periplasmic WT β-lactamase, fusion of β-lactamase to the GlyGly-CTERM of VesB (β-lac-GG) resulted in β-lactamase activity in both the periplasmic and membrane fractions. Fusion of the IG fold from VesB onto β-lactamase (β-lac-IG) did not shift its subcellular location but resulted in an increase in activity relative to WT β-lac in the periplasmic fraction, potentially the result of increased stability of this construct. We did not observe a significantly higher enzyme activity in the culture supernatants of β-lac-GG or β-lac-IG as compared to WT β-lac, indicating that both the IG fold and GlyGly-CTERM are required to facilitate transport to the extracellular environment. To determine if any of the β-lactamase constructs were localized to the cell surface, we quantified the β-lactamase activity of intact cells and sonicated cells using the substrate nitrocefin (**Figure 5C**). Nitrocefin is not readily permeable across the gram-negative cell envelope and has been used previously to detect defects in outer membrane integrity (20). We reasoned that the activity associated with intact cells reflects mostly surface exposed β-lactamase. When we assessed the enzyme activity of intact cells expressing WT β-lactamase, β-lac-GG or β-lac-IG, we observed only a small percentage (< 20%) as compared to the sonicated cells suggesting that the majority of these chimeras remained intracellular. The activity associated with the same constructs when expressed in the Δ*epsD* mutant strain was indistinguishable from the WT strain, indicating limited surface localization of these constructs. In contrast, WT cells expressing β-lac-IG/GG had significantly higher surface activity (65%) that was reduced to 25% for the Δ*epsD* mutant strain. Collectively, these results indicate that the GlyGly-CTERM positions β-lactamase in the inner membrane where it is processed by the rhomboid proteases. An additional secretion signal, in this case the IG fold from VesB, is required for outer membrane translocation via the T2SS.

## Discussion

Previously we have demonstrated that VesB is intimately connected to RssP via its GlyGly-CTERM domain. Using chimeric proteins, we demonstrate here that three additional GlyGly-CTERM motifs from VesA, VesC, and VC1485 are similarly recognized by RssP, resulting in efficient activation of VesB. Subsequent proteomic analysis suggested that VC1485, VesA, and perhaps Xds are targeted to the cell membrane fraction, likely the cell surface, in an RssP dependent manner. Our findings for Xds are corroborated by a previous report indicating that Xds is extracellular but associated with the cell (21). Surprisingly, we found that VesC, despite its GlyGly-CTERM being able to functionally replace that of VesB to promote the biogenesis of active VesB, is primarily found in the culture supernatant. We demonstrated that this is due to proteolytic processing at an intrinsic site in the unstructured linker N-terminal to the GlyGly-CTERM of VesC (**Figure 6A**). The equivalent region in VesB is subject to proteolysis by the rhomboid protease GlpG prior to outer membrane translocation when RssP is absent (12). In addition, when VesB was previously purified from *V. cholerae* culture supernatants there was evidence for a ‘ragged C-terminus’ of VesB potentially due to additional non-specific proteolysis of the unstructured region (6). Despite the release of VesC to the culture supernatant, chimeric constructs clearly indicate that the GlyGly-CTERM of VesC is still preferentially targeted to RssP (**Figure 6B**) and not GlpG, and collectively, the various GlyGly-CTERM motifs found in *V. cholerae* target their passenger proteins to RssP for processing. Interestingly, in the absence of RssP, VesB and perhaps VC1485 seem to be released to the culture supernatant by GlpG although it is unclear at what site VC1485 might be cleaved by GlpG.

Previously, we determined that expression with the GlyGly-CTERM is required for the eventual processing by RssP and surface localization of VesB (12) and that when coupled with the IG fold from VesB it is sufficient to direct β-lactamase for full secretion (9). In this previous study, a significant portion of the enzyme activity of the β-lac-IG/GG construct was associated with the cell pellet (9), which we show here is partially contributed by surface localized β-lac-IG/GG. Combined with the data on the HapA constructs, we further show that the GlyGly-CTERM from VesB works in concert with a secretion signal, the IG fold from VesB or a yet to be identified secretion signal in HapA, to direct passenger proteins to the cell surface once processed by RssP. While not shown directly, we speculate that membrane retention via the GlyGly-CTERM motif initially occurs at the inner membrane during signal peptide mediated transport and prior to outer membrane translocation by the T2SS (9). Combined with the observation that β-lac-IG is primarily found in the periplasmic fraction of cells and not secreted, we pose that inner membrane attachment improves the efficiency of secretion of proteins with a GlyGly-CTERM motif. Secretion from an inner membrane location is not unique to GlyGly-CTERM proteins, as *A. baumannii* secretes an effector that is initially associated with a membrane contained chaperone (22). Similarly, it has been suggested that prior binding of the lipoprotein PulA to the inner membrane supports its interaction with the T2SS in *K. pneumoniae* (23). We also note that homologs of GlyGly-CTERM proteins have been bioinformatically identified with other, distinct sorting motifs or with no apparent sorting motif supporting our findings that the GlyGly-CTERM is a discrete motif that functions independently of the passenger protein akin to N-terminal signal peptides (11, 24). In this model, the GlyGly-CTERM targets passenger proteins to RssP resulting in their posttranslational modification and inner membrane retention where they can be recognized by the T2SS for outer membrane translocation.

Regulation of proteases and extensive posttranslational modifications have been described for HapA as well as the serine protease IvaP both expressed and secreted in a T2SS dependent manner by *V. cholerae* (19, 25). We propose that the GlyGly-CTERM functions, in part, to regulate its secretion to prevent off-target intracellular activity. This is supported by previous findings showing that inactivating mutations arise within the coding region of VesC in T2SS mutant strains, suggesting a detrimental effect of VesC periplasmic accumulation (17). Previously, overexpression of VesC in T2SS mutant strains were shown to prevent growth (17). Most GlyGly-CTERM proteins are predicted to harbor hydrolytic activities (11) and it seems logical that *V. cholerae* would benefit from keeping these enzymes attached to the inner membrane (in inactive forms) prior to transport across the outer membrane to prevent off-target effects if freely soluble in the periplasmic space. Perhaps the presence of multiple enzymes freely released to the periplasm would be especially detrimental to the cell.

Overall, the data presented here further our understanding of how *V. cholerae* regulates the secretion and surface localization of various hydrolytic enzymes expressed with a GlyGly-CTERM extension. Numerous other bacteria harbor GlyGly-CTERM-containing proteins including pathogenic species such as *Acinetobacter baumannii* and *Aeromonas hydrophilia* as well as non-pathogenic environmental species such as *Shewanella oneidensis* (11) In addition, unique C-terminal sorting motifs have been described in gram-negative bacteria such as *Myxobacteria* and it is believed that proteins with these motifs also localize to the cell surface in a T2SS dependent manner (26). Other, thus far, uncharacterized sorting motifs have been identified bioinformatically in diverse bacteria warranting further study (24).

## Materials and Methods

### Bacteria strains, plasmids, and Growth conditions

The *V. cholerae* strain El Tor O1 strains, N16961, and the previously constructed isogenic mutants NΔ*vesABC*, NΔ*glpG*, N*rssP*::kan, NΔ*epsD*, NΔ*glpG rssP*::kan, were used in this study (6, 27). All plasmids and primers used in this study are listed in Table S1. All polymerase chain reactions (PCR), cloning, and restriction enzyme digestions were done with either SuperFi Platinum polymerase or PfuTurbo, T4 DNA ligase, and restriction enzymes from New England Biolabs and primers that were synthesized at IDT Technologies. pCR-Script (Stratagene) and pMMB67EH constructs were transformed into *E. coli* MC1061 and pCVD442 constructs into SY327λpir. Triparental conjugation was performed with a helper strain carrying pRK2013 to transfer plasmids into N16961 and its isogenic mutants. Overhanging primers coding for the GlyGly-CTERM of VesA, VesC and VC1485 were used to make the VesB chimeric proteins. Site-specific mutants were generated using overlapping primers and PfuTurbo followed by DpnI digestion and transformation into MC1061 per manufacturer’s instruction. IPTG was added for the induction of expression where indicated. For experiments expressing β-lactamase or chimeric constructs, a derivative of pMMB67EH, pMMB960, containing a kanamycin cassette for antibiotic selection was used (28). Kanamycin was supplemented at 50 µg/mL.

### Subcellular fractionation and outer membrane vesicle isolation

Cell pellets were fractionated as described previously (29). To assess membrane content, an NADH dehydrogenase assay was used as described with NADH as substrate by monitoring the loss of optical density at 340 nm (30). Percent activity was similarly calculated by dividing each individual value by the total activity across each sample.

For isolation of crude outer membrane vesicles (OMVs), cells were grown overnight in LB supplemented with 10 µM IPTG before culture supernatants were isolated by centrifugation at 10,000 × g for 10 minutes. Subsequent sterile filtration and centrifugation at 200,000 × g for 3 h pelleted the insoluble fraction. Separately, the cleared supernatant and the pellets containing crude outer membrane vesicles resuspended in LB were assayed for protease activity using the fluorogenic peptide N-tert-butoxycarbonyl-Gln-Ala-Arg-7-amido-4-methylcoumarin.

### Sodium dodecyl sulfate-polyacrylamide gel electrophoresis, immunoblotting, serine hydrolase labeling, zymography, HapA milk agar spot assay

Samples were prepared and analyzed by sodium dodecyl sulfate-polyacrylamide gel electrophoresis and immunoblotting as described previously. Monoclonal antiserum against His (Fisher) was incubated with the nitrocellulose membrane for 2 h (1:5,000). Horseradish peroxidase-conjugated goat anti-rabbit immunoglobulin G (Bio-Rad) used at 1:20,000 was incubated with the membrane for 1 h. Membranes were developed using ECL 2 Western blotting reagent (Fisher) and visualized using a Typhoon V variable mode imager system and ImageJ imager software. For serine hydrolase labeling, cell, supernatant, cytoplasmic, periplasmic, or membrane fractions were incubated with Activ TAMRA-FP Serine Hydrolase Probe (Fisher) for 2 hours in the dark before performing SDS-PAGE. Gels were visualized with fluorescent gel imaging using a Typhoon V variable mode imager system and ImageJ imager software. For zymogram gels, without heating samples were run on Novex 10% Zymogram gels containing gelatin (Fisher). Gels were renatured, developed, and stained with Coomassie blue for visualization according to manufactures recommendations. For HapA spot assays, overnight cultures of *V. cholerae* strains were spotted onto LB agar containing 1% skim milk and incubated overnight at 37 ° C.

### Protease assay

*V. cholerae* supernatants and intact cells were measured for VesB and VesC protease activity using N-tert-butoxycarbonyl-Gln-Ala-Arg-7-amido-4-methylcoumarin as described previously (9). For detection of VesA specific proteolytic activity, we used the fluorogenic peptide Suc-LeuLeu-Val-Tyr-7-amino-4-methylcoumarin. Change in fluorescence per minute was calculated and converted to moles of methylcoumarin (AMC) generated per minute via a standard curve with known concentrations of AMC. The rate of AMC generation was normalized by OD_600_ of the cultures. For time course experiments, protease activity was assessed as described at the indicated time points. Activity represents a percentage of the total activity across the cell and supernatant fraction at each time point.

### Beta-lactamase

Beta-lactamase activity was assessed by analyzing the hydrolysis of nitrocefin and presented as arbitrary units as previously described (9). Briefly, nitrocefin resuspended in PBS, and 5% DMSO was added to isolated cells or culture supernatants to a final concentration of 100 µM and the absorbance at 500 nm was recorded using an H1 Synergy plate reader. Sonication-disrupted cells or intact cells were tested using the same protocol.

### Cell surface detection of HapA

Cells were washed, blocked with 2% BSA, and incubated with 1:1,000 of HapA antiserum. Following incubation with 1:1,000 of Alexa Fluor 488 F(ab′)2 goat anti-rabbit immunoglobulin G (Invitrogen) and washing, fluorescence was measured (Ex 488 nm/Em 525 nm). The results were normalized to the fluorescence intensity of the cell culture of the same strain carrying an empty vector as described (9). The cells were also visualized by differential interference contrast and fluorescent microscopy using a Nikon Eclipse 90i fluorescence microscope equipped with a Nikon Plan Apo VC 100× (1.4 numerical aperture) oil immersion objective and a CoolSNAPHQ2 digital camera as previously described (6).

### Protein precipitation and LC-MS/MS of cell membrane and culture supernatants

Wild-type (WT) *V. cholerae* N16961, *rssP*::kan, or Δ*glpG* mutants were grown at 37°C for 16 h in Luria-Bertani broth (LB). Culture supernatants were isolated via centrifugation and filter sterilized before being subjected to protein precipitation with an equal volume of pyrogallol red-molybdate-methanol (PRMM) solution as previously described (9). Normalized samples were subjected to GeLC10-tandem mass spectrometry (LC-MS/MS) and label-free quantitation as previously described (9). Normalized Spectral Abundance Factor (NSAF) was used to compare three biological samples between the different strains. Scaffold 5 software was used to collate, filter, and analyze the data. The complete mass spectrometry data set (PXD050626) was deposited to the ProteomeXchange Consortium (http://proteomecentral.proteomexchange.org). Insoluble (membrane) cell material was isolated as previously described and subjected to the same analysis (PXD050605) (31).

### Statistical analysis

All data are presented as mean ± standard deviation. The enzyme assays and growth curves were performed with ≥3 biological replicas in technical triplicates. Differences between two groups in the LC-MS/MS analysis were determined by a paired, two-tailed Student’s *t*-test. ANOVA with a Dunnett correction for multiple comparisons was used to compare three or more mutants to specified controls. Tukey or Sidak’s correction was used for multiple comparisons of three or more samples where every mean was compared to every other mean. Results yielding a *P* value of <0.05 were considered statistically significant, and only significant differences are indicated. All calculations were done using GraphPad Prism version 10.0.0 for Windows, GraphPad Software, Boston, MA, USA; https://www.graphpad.com).

### Structural prediction

Structural models for the interaction of VesB chimeras and RssP were generated from primary amino acid sequence using AlphaFold3 (32). Structures were visualized with ChimeraX (33).

## Acknowledgements

The anti-HA/protease antibodies and Δ*hapA* mutant strain were kind gifts of Dr. Richard Finkelstein. This work was, in part, supported by R01AI137085 (to M.S.) and F32AI178852 (to C.R.).

## References

1. Montero DA, Vidal RM, Velasco J, George S, Lucero Y, Gomez LA, Carreno LJ, Garcia-Betancourt R, O’Ryan M. 2023. Vibrio cholerae, classification, pathogenesis, immune response, and trends in vaccine development. Front Med (Lausanne) 10:1155751.

2. Korotkov KV, Sandkvist M. 2019. Architecture, Function, and Substrates of the Type II Secretion System. EcoSal Plus 8.

3. Baryalai P, Irenaeus D, Toh E, Ramstedt M, Uhlin BE, Nadeem A, Wai SN. 2025. Hemagglutinin Protease HapA Associated With Vibrio cholerae Outer Membrane Vesicles (OMVs) Disrupts Tight and Adherens Junctions. J Extracell Vesicles 14:e70092.

4. d’Enfert C, Chapon C, Pugsley AP. 1987. Export and secretion of the lipoprotein pullulanase by Klebsiella pneumoniae. Mol Microbiol 1:107–16.

5. Ferrandez Y, Condemine G. 2008. Novel mechanism of outer membrane targeting of proteins in Gram-negative bacteria. Mol Microbiol 69:1349–57.

6. Gadwal S, Johnson TL, Remmer H, Sandkvist M. 2018. C-terminal processing of GlyGly-CTERM containing proteins by rhombosortase in Vibrio cholerae. PLoS Pathog 14:e1007341.

7. Adams CO, Campbell JA, Zhang B, Cleaver L, Bier SB, Mayoral J, White RC, Garnett JA, Cianciotto NP. 2025. Legionella pneumophila type II secretome reveals a polysaccharide deacetylase that impacts intracellular infection, biofilm formation, and resistance to polymyxin-and serum-mediated killing. mBio 16:e0139325.

8. Jackson-Litteken CD, Di Venanzio G, Le NH, Scott NE, Djahanschiri B, Distel JS, Pardue EJ, Ebersberger I, Feldman MF. 2022. InvL, an Invasin-Like Adhesin, Is a Type II Secretion System Substrate Required for Acinetobacter baumannii Uropathogenesis. mBio 13:e0025822.

9. Shannon A, Johnson T, Roberts CS, Chaton CT, Korotkov KV, Sandkvist M. 2025. The PDZ domain of EpsC is required for extracellular secretion of VesB by the Type II secretion system in Vibrio cholerae. J Bacteriol doi: 10.1128/jb.00144-25:e0014425.

10. Francetic O, Pugsley AP. 2005. Towards the identification of type II secretion signals in a nonacylated variant of pullulanase from Klebsiella oxytoca. J Bacteriol 187:7045–55.

11. Haft DH, Varghese N. 2011. GlyGly-CTERM and rhombosortase: a C-terminal protein processing signal in a many-to-one pairing with a rhomboid family intramembrane serine protease. PLoS One 6:e28886.

12. Roberts CS, Shannon AB, Korotkov KV, Sandkvist M. 2024. Differential processing of VesB by two rhomboid proteases in Vibrio cholerae. mBio 15:e0127024.

13. Sikora AE, Zielke RA, Lawrence DA, Andrews PC, Sandkvist M. 2011. Proteomic analysis of the Vibrio cholerae type II secretome reveals new proteins, including three related serine proteases. J Biol Chem 286:16555–66.

14. Gadwal S, Korotkov KV, Delarosa JR, Hol WG, Sandkvist M. 2014. Functional and structural characterization of Vibrio cholerae extracellular serine protease B, VesB. J Biol Chem 289:8288–98.

15. Mondal A, Tapader R, Chatterjee NS, Ghosh A, Sinha R, Koley H, Saha DR, Chakrabarti MK, Wai SN, Pal A. 2016. Cytotoxic and Inflammatory Responses Induced by Outer Membrane Vesicle-Associated Biologically Active Proteases from Vibrio cholerae. Infect Immun 84:1478–1490.

16. Seper A, Hosseinzadeh A, Gorkiewicz G, Lichtenegger S, Roier S, Leitner DR, Rohm M, Grutsch A, Reidl J, Urban CF, Schild S. 2013. Vibrio cholerae evades neutrophil extracellular traps by the activity of two extracellular nucleases. PLoS Pathog 9:e1003614.

17. Rule CS, Park YJ, Delarosa JR, Turley S, Hol WGJ, McColm S, Gura C, DiMaio F, Korotkov KV, Sandkvist M. 2020. Suppressor Mutations in Type II Secretion Mutants of Vibrio cholerae: Inactivation of the VesC Protease. mSphere 5.

18. Meibom KL, Blokesch M, Dolganov NA, Wu CY, Schoolnik GK. 2005. Chitin induces natural competence in Vibrio cholerae. Science 310:1824–7.

19. Benitez JA, Silva AJ. 2016. Vibrio cholerae hemagglutinin(HA)/protease: An extracellular metalloprotease with multiple pathogenic activities. Toxicon 115:55–62.

20. Hancock RE, Wong PG. 1984. Compounds which increase the permeability of the Pseudomonas aeruginosa outer membrane. Antimicrob Agents Chemother 26:48–52.

21. Pressler K, Mitterer F, Vorkapic D, Reidl J, Oberer M, Schild S. 2019. Characterization of Vibrio cholerae’s Extracellular Nuclease Xds. Front Microbiol 10:2057.

22. Kinsella RL, Lopez J, Palmer LD, Salinas ND, Skaar EP, Tolia NH, Feldman MF. 2017. Defining the interaction of the protease CpaA with its type II secretion chaperone CpaB and its contribution to virulence in Acinetobacter species. J Biol Chem 292:19628–19638.

23. East A, Mechaly AE, Huysmans GHM, Bernarde C, Tello-Manigne D, Nadeau N, Pugsley AP, Buschiazzo A, Alzari PM, Bond PJ, Francetic O. 2016. Structural Basis of Pullulanase Membrane Binding and Secretion Revealed by X-Ray Crystallography, Molecular Dynamics and Biochemical Analysis. Structure 24:92–104.

24. Haft DH. 2024. In silico discovery of the myxosortases that process MYXO-CTERM and three novel prokaryotic C-terminal protein-sorting signals that share invariant Cys residues. J Bacteriol 206:e0017323.

25. Howell M, Dumitrescu DG, Blankenship LR, Herkert D, Hatzios SK. 2019. Functional characterization of a subtilisin-like serine protease from Vibrio cholerae. J Biol Chem 294:9888–9900.

26. Guo T, Haft DH, Wall D. 2025. Myxosortase: an intramembrane protease that sorts MYXO-CTERM proteins to the cell surface. mBio 16:e0406724.

27. Sikora AE, Lybarger SR, Sandkvist M. 2007. Compromised outer membrane integrity in Vibrio cholerae Type II secretion mutants. J Bacteriol 189:8484–95.

28. Waack U, Warnock M, Yee A, Huttinger Z, Smith S, Kumar A, Deroux A, Ginsburg D, Mobley HLT, Lawrence DA, Sandkvist M. 2018. CpaA Is a Glycan-Specific Adamalysin-like Protease Secreted by Acinetobacter baumannii That Inactivates Coagulation Factor XII. mBio 9.

29. Sandkvist M, Hough LP, Bagdasarian MM, Bagdasarian M. 1999. Direct interaction of the EpsL and EpsM proteins of the general secretion apparatus in Vibrio cholerae. J Bacteriol 181:3129–35.

30. Cian MB, Giordano NP, Mettlach JA, Minor KE, Dalebroux ZD. 2020. Separation of the Cell Envelope for Gram-negative Bacteria into Inner and Outer Membrane Fractions with Technical Adjustments for Acinetobacter baumannii. J Vis Exp doi: 10.3791/60517.

31. Roberts CS, Ni F, Mitra B. 2021. The Zinc and Iron Binuclear Transport Center of ZupT, a ZIP Transporter from Escherichia coli. Biochemistry 60:3738–3752.

32. Abramson J, Adler J, Dunger J, Evans R, Green T, Pritzel A, Ronneberger O, Willmore L, Ballard AJ, Bambrick J, Bodenstein SW, Evans DA, Hung CC, O’Neill M, Reiman D, Tunyasuvunakool K, Wu Z, Zemgulyte A, Arvaniti E, Beattie C, Bertolli O, Bridgland A, Cherepanov A, Congreve M, Cowen-Rivers AI, Cowie A, Figurnov M, Fuchs FB, Gladman H, Jain R, Khan YA, Low CMR, Perlin K, Potapenko A, Savy P, Singh S, Stecula A, Thillaisundaram A, Tong C, Yakneen S, Zhong ED, Zielinski M, Zidek A, Bapst V, Kohli P, Jaderberg M, Hassabis D, Jumper JM. 2024. Accurate structure prediction of biomolecular interactions with AlphaFold 3. Nature 630:493–500.

33. Meng EC, Goddard TD, Pettersen EF, Couch GS, Pearson ZJ, Morris JH, Ferrin TE. 2023. UCSF ChimeraX: Tools for structure building and analysis. Protein Sci 32:e4792.

