## Supplemental Figures for "The GlyGly-CTERM domain functions as an independent motif that targets proteins to rhombosortase in *Vibrio cholerae*"

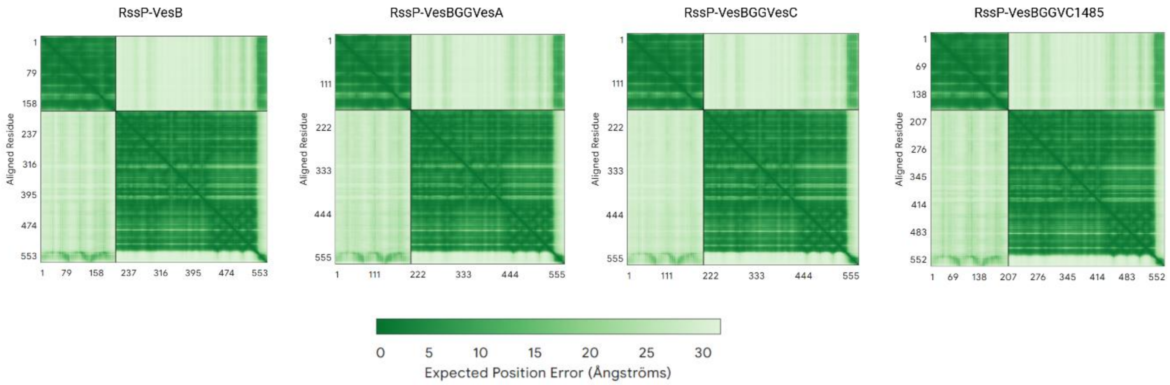

**Supplemental Figure 1. Alignment error for RssP and VesB chimeric protein interaction.** The primary amino acid sequences of RssP and VesB or VesB chimera were used to predict the interaction using AlphaFold3. Position error shown.

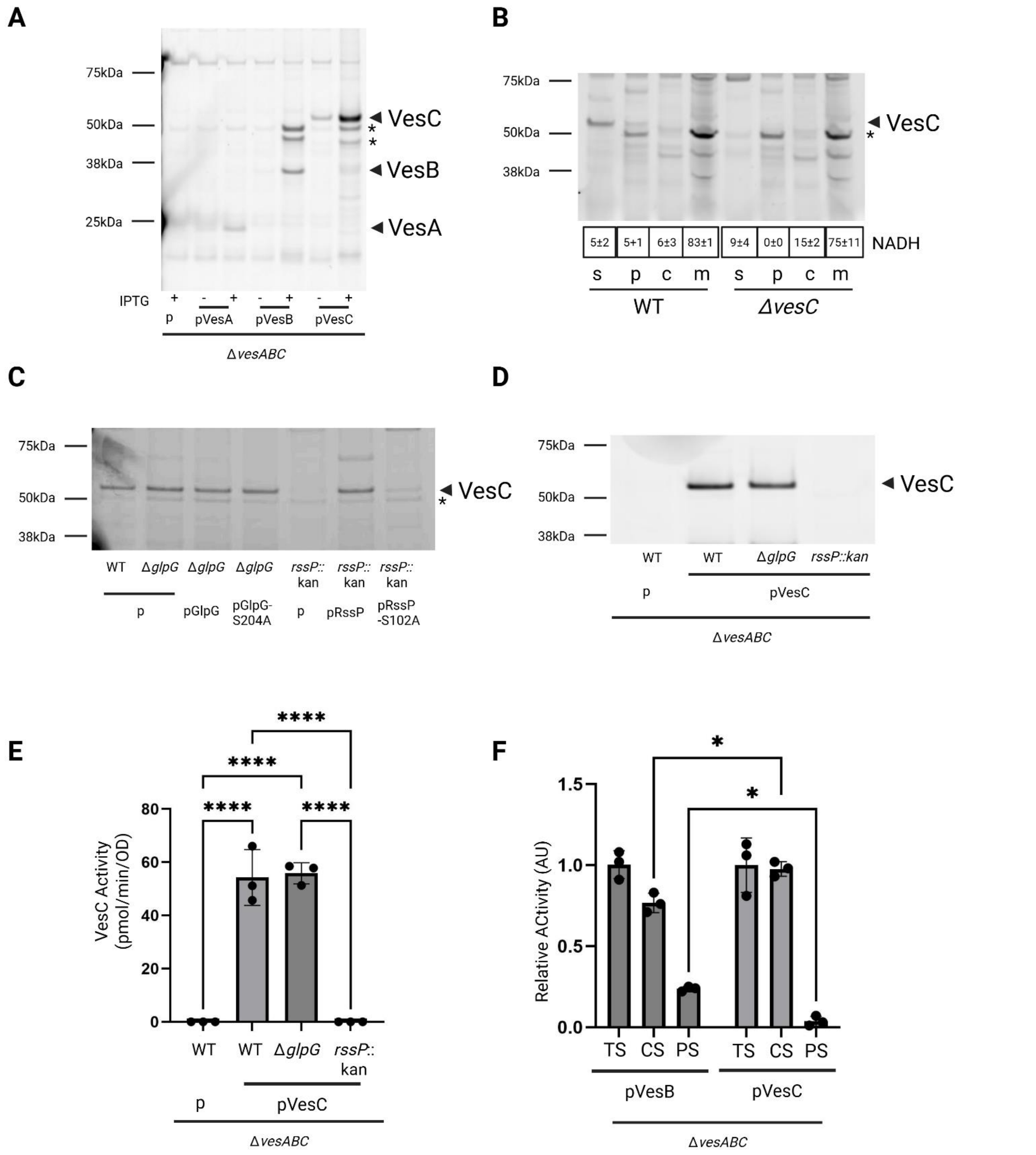

**Sup. Figure 2. VesC is secreted into the extracellular space in an RssP-dependent manner. A.** Supernatants from overnight cultures of WT strain N16961 containing empty vector or plasmids expressing indicated Ves protease were isolated, incubated with ActivX serine hydrolase probe, before separation by SDS-PAGE and fluorescent gel imaging. Where indicated, IPTG was used at 50 µM for expression of VesA and VesB and 10 µM for VesC. VesA, VesB, and VesC and molecular weight markers are indicated. Representative gel is shown (n=2). Asterisks show non-specific bands. **B.** Log phase cultures of WT or Δ*vesC* mutant cells were fractionated into supernatant (s), periplasmic (p), cytoplasmic (c), and membrane (m) fractions, incubated with ActivX serine hydrolase probe, before separation by SDS-PAGE and fluorescent gel imaging. As a control for membrane isolation, NADH dehydrogenase activity was monitored by measuring decrease in absorbance at 340 nm over time. Average ± standard deviation from technical duplicates is shown. Molecular weight markers and VesC are indicated. Non-specific band is indicated with asterisk. Representative gel is shown (n=2). **C.** Supernatants from overnight cultures of WT and indicated rhomboid mutants containing empty vector or complemented with indicated plasmid were isolated, incubated with ActivX serine hydrolase probe before separation by SDS-PAGE and fluorescent gel imaging. VesC and molecular weight markers are indicated. Representative gel is shown (n=2). **D.** Supernatants from WT and indicated rhomboid mutants containing empty vector or ectopically overexpressing (10 μM IPTG) VesC were processed and visualized as in **C**. VesC and molecular weight markers are indicated. Representative gel is shown (n=2). **E.** Supernatants from **D** were assessed for serine protease activity against the fluorogenic peptide Boc-Gln-Ala-Arg-AMC. Data represent mean ± SD of $n$ = 3 experiments in technical triplicate with Tukeys multiple comparison test with **** representing $p \le 0.0001$. Only statistically significant comparisons are shown. **F.** Supernatants from overnight cultures of ΔvesABC mutant strain ectopically overexpressing (10 μM IPTG) VesB or VesC were isolated (S), separated into cleared supernatant (CS) and pelleted supernatant (PS) before assessing for serine protease activity against the fluorogenic peptide Boc-Gln-Ala-Arg-AMC. Activity is expressed relative to the total supernatant (S) activity. Data represent mean ± SD of $n$ = 3 experiments in technical triplicate with Tukeys multiple comparison test with * representing $p<0.05$.

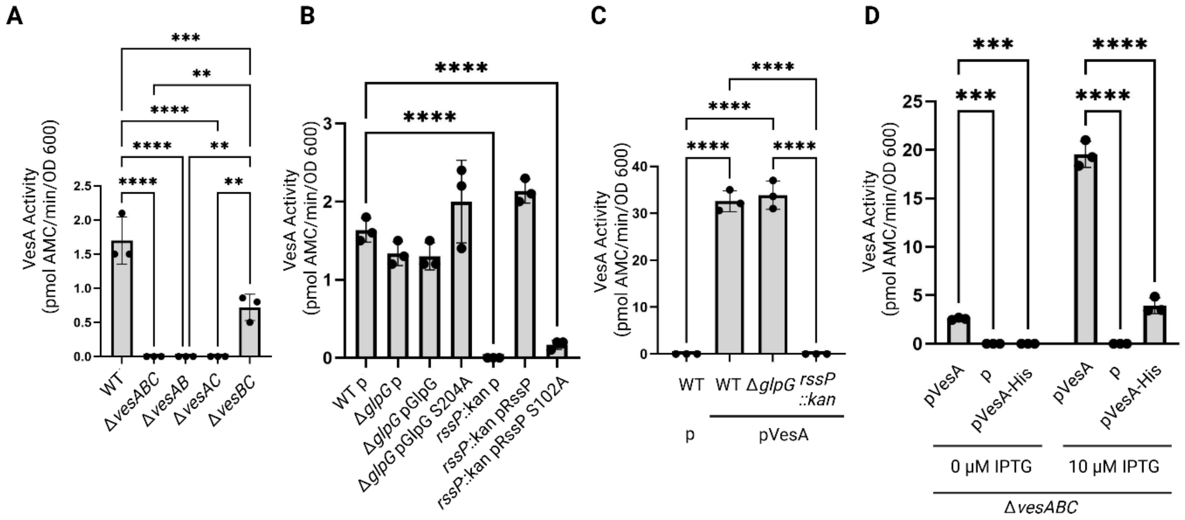

**Sup. Figure 3. VesA requires RssP and an intact GlyGly-CTERM for efficient activation. A.** Supernatants from overnight cultures of WT and indicated mutant strains were isolated and assessed for serine protease activity using the fluorogenic peptide substrate Suc-Leu-Leu-Val-Tyr-AMC. Data represent mean ± SD of *n* = 3 experiments in technical triplicate with Tukeys multiple comparison test with ** representing p<0.005, *** p<0.0005 and **** p≤ 0.0001. Only significant differences are shown. **B.** Supernatants from overnight cultures of WT and indicated rhomboid mutant strains ectopically complemented with WT rhomboid protease or catalytic mutant were isolated and assessed for serine protease activity using the fluorogenic peptide substrate Suc-Leu-Leu-Val-Tyr-AMC. Data represent mean ± SD of *n* = 3 experiments in technical triplicate with Dunnett's multiple comparison test with **** representing p≤0.0001. Only significant differences are shown. **C.** Supernatants from overnight cultures of WT and indicated mutant strain containing empty vector (p) or overexpressing (50 µM IPTG) plasmids encoding WT VesA were isolated and assessed for serine protease activity using the fluorogenic peptide substrate Suc-Leu-Leu-Val-Tyr-AMC. Data represents ± SD of *n* = 3 experiments in technical triplicate with Tukeys multiple comparison test with *** representing p<0.0005 and **** p≤0.0001. Only significant differences are shown. **D.** Supernatants from overnight cultures of the mutant strain Δ*vesABC* containing empty vector (p) or plasmids encoding WT VesA or indicated VesA mutant supplemented with indicated IPTG concentration were isolated and assessed for serine protease activity using the fluorogenic peptide substrate Suc-Leu-Leu-Val-Tyr-AMC. Data represent mean ± SD of *n* = 3 experiments in technical triplicate with Sidak's multiple comparison test with *** representing p<0.0005 and **** p≤ 0.0001. Only significant differences are shown.

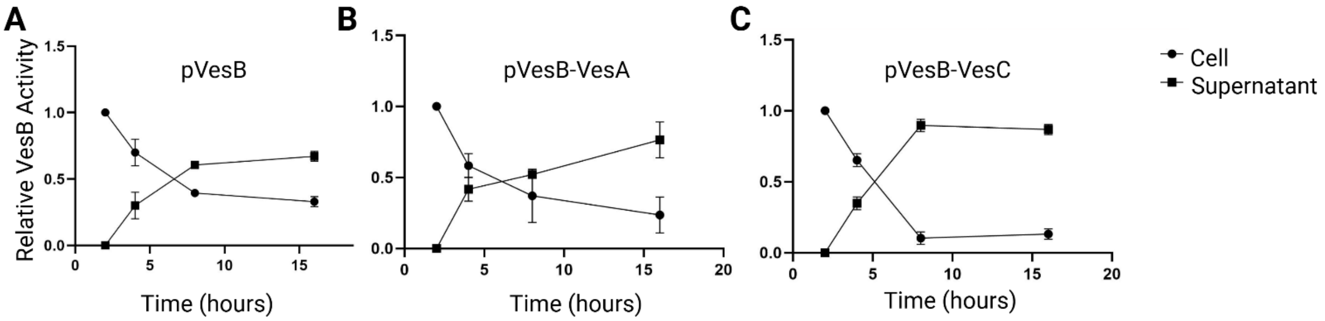

**Sup. Figure 4. The GlyGly-CTERM and disordered region from VesC transiently associates the passenger protein with the cell fraction.** Mutant strain Δ*vesABC* expressing VesB (**A**), VesB-VesA (**B**), or VesB-VesC (**C**) chimera were grown for 16 hours. At indicated timepoints, samples were taken and separated into cell and supernatant fractions before they were assessed for serine protease activity against the fluorogenic peptide Boc-Gln-Ala-Arg-AMC. Relative activity was calculated by taking the cell and supernatant activity, respectively, and diving by the total activity across the cell and supernatant fractions at the indicated time point. Data represent mean ± SD of *n* = 3 experiments in technical triplicate.

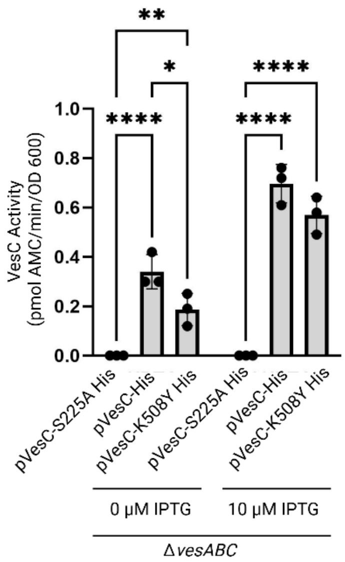

**Sup. Figure 5. VesC-K508Y His6 is active.** Culture supernatants from the Δ*vesABC* mutant strain ectopically expressing different VesC-His6 variants in the absence and presence of IPTG were isolated and assessed for serine protease activity against the fluorogenic peptide Boc-Gln-Ala-Arg-AMC. Data represent mean ± SD of *n* = 3 experiments in technical triplicate with Sidak post hoc analysis with * representing p≤0.05 **p≤0.005, and ****p≤0.0001. Only statistically significant comparisons are shown.

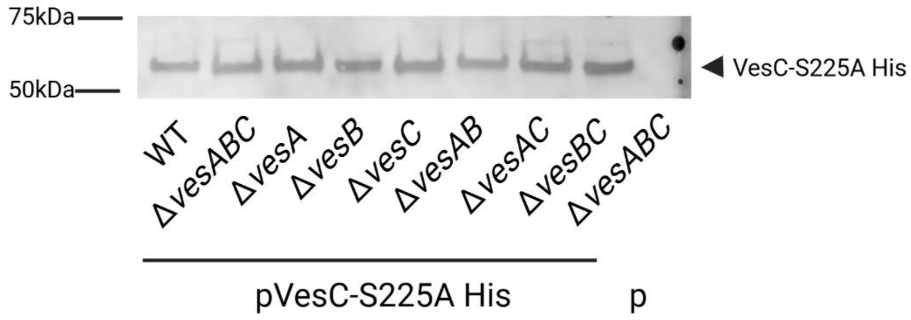

**Sup. Figure 6. VesC-His is not processed in trans.** Culture supernatants from indicated mutant strains containing empty vector or encoding catalytically inactive VesC-His6 were isolated, run on SDS-PAGE, transferred to nitrocellulose, and blotted against anti-His6 antibodies. VesC species and molecular weight markers are indicated. Representative blot is shown (n=2).

### Tables S1. Strains and plasmids used in this study.

| Plasmid/strain | Description | Fwd Primer 5' to 3' | Rev Primer 5' to 3' |
|---|---|---|---|
| N16961 | El tor, *Vibrio cholerae* strain | | |
| NΔ*glpG* (1) | in frame deletion of *glpG* in N16961 | | |
| NΔ*epsD* (2) | in frame replacement of *epsD* with kanamycin cassette | | |
| N *rssP*::kan (1) | Insertion of kanamycin cassette into *rssP* | | |
| NΔ*glpG* *rssP*::kan (1) | *glpG* and *rssP* inactivated | | |
| NΔ*hapA* | Kind gift from Dr. Richard Finkelstein | | |
| NΔ*vesABC* (3) | *vesA, B, C* mutant | | |
| NΔ*vesAB* (3) | *vesA, B* mutant | | |
| NΔ*vesBC* (3) | *vesB, C* mutant | | |
| NΔ*vesAC* (3) | *vesA, C* mutant | | |
| NΔ*vesC* (3) | in frame deletion of *vesC* | | |
| NΔ*vesA* (3) | In frame replacement of *vesA* with Cm cassette | | |
| NΔ*vesB* (3) | In frame replacement of *vesB* with Kan cassette | | |
| pRK2013 (4) | Helper Strain | | |
| SY327λpir (5) | Helper strain | | |
| pMMB67 (7) | Expression vector (7) | | |
| pMMB960 (8) | pMMB67 with $Kan^r$ instead of $Amp^r$ | | |
| pCVD442 (5) | Suicide vector (5) | | |
| pRssP (1) | RssP expressed from pMMB67 | | |
| pRssP S102A (1) | Catalytic serine mutated to alanine | | |
| pGlpG (9) | GlpG expressed from pMMB67 | | |
| pGlpG S204A | Catalytic serine | | |

| | | | |
|---|---|---|---|
| | mutated to alanine | | |
| pHapA (10) | HapA expression from pMMB66EH | | |
| pHapA-GG | Used pHapA and pVesB as template | GGCAAAAGGCGAGTCCAAGCGTAACGTCACACCAGA | TTACGCTTGGACTCGCCTTTTGCCCCAAGTGCCTCT |
| pβ-lac (2) | B-lactamase gene from pMMB67EH | | |
| pβ-lac-IG | Used pβ-lac and GlyGly-CTERM VesB mutant as template, pMMB960 | CTCACTGATTAAGCATTGGCACACTAACCAGCTTAGCTATGATC | GCTAAGCTGGTTAGTGTGCCAATGCTTAATCAGTGAGGC |
| pβ-lac-GG | Used pβ-lac and WT pVesB as template, pMMB960 | TCACTGATTAAGCATTGGTCGCCTTTTGCCCCAAG | GGGCAAAAGGCGACCAATGCTTAATCAGTGAGGC |
| pβ-lac-IG/GG (2) | Used β-lac and WT VesB as template, pMMB960 | | |
| pVesB (3) | WT VesB | CTCACTGATTAAGCATTGGCACACTAACCAGCTTAGCTATGATC | GCTAAGCTGGTTAGTGTGCCAATGCTTAATCAGTGAGGC |
| pVesBGGVesA | Used pVesB and pVesA as template | GGAGGAGGTGTCTCTTTGCTTATTGCTTTC | ACCTCCTCCAGAAGAGGCACTTGG |
| pVesBGGVesC | Used pVesB and pVesC as template | ACCTAGGCTACCACCGCCAGAAGAGGCACTTGGCGA | TCTTCTGGCGGTGGTAGCCTAGGTGGGGCAGCCTTA |
| pVesBGGVC1485 | Used pVesB and pVC1485 as template | GGGGGTAGCTTCGGATTAGGG | GCTACCCCCAGAAGAGGCACT |
| pVC1485 | Used N16961 as template, VC1485 cloned into pMMB67 | CAGCGGATCCACAGGAAAGTGAATAGAA | TTATCTGCAGTTGTGCCCTTCCTGACA |
| pVesB-VesA | Used WT pVesA and pVesB as template | TCGTATTCAACTGGATACTGGAGGGACTACAACGGTTAGC | AACCGTTGTAGTCCTCCAGTATCCAGTTGAATACGAGACGT |
| pVesB-VesC | Used pVesA and pVesB as template | TCGTATTCAACTGGATACTCCAAAATTCTTTGATCTACCTCCA | AGGTAGATCAAAGAATTTTGGAGTATCCAGTTGAATACGAGACGT |
| pVesC (3) | WT VesC | | |
| pVesC-K508Y | Point mutant | GCCAAATAAGCTACGATCCATATTTCTTTGATCTACCTCCACC | GGTGGAGGTAGATCAAAGAAATATGGATCGTAGCTTATTTGGC |
| pVesC-R519Y | Point mutant | GATCTACCTCCACCGCTTGATATTTATATCGGTGATAGTGG | CCACTATCACCGATATAAATATCAAGCGGTGGAGGTAGATC |
| pVesAΔ23-His (2) | Used pVesA as template, phosphorylated primers | CATCACCATTGAACTTTATGATCAAGGTGTG | GTGATGGTGTCCTCCAGATGAGCTCTC |
| pVesAΔ23-His S221A (2) | Previously generated | | |
| pVesBΔ20-His (2) | Used pVesB as template | CATCATCACCATCATCACTGATTACCTATCCGAGATCTG | GTGATGATGGTGATGATGACCGCCAGAAGAGGCACTTG |
| pVesBΔ20-His S221A (2) | Previously generated | | |
| pVesCΔ22-His (2) | Used pVesB as template | CATCATCACTGATCTATCGATAGCCAGCATGC | GTGATGATGACCGCCGCCACTATC |
| pVesCΔ22-His S225A (2) | Previously generated | | |
| pVesCΔ22-His K508Y | Used K508Y | | |

| | |
|---|---|
| | primers with pVesC Δ22-His as template |
| pVesA (3) | WT VesA |

Supplementary References

1. Gadwal S, Johnson TL, Remmer H, Sandkvist M. 2018. C-terminal processing of GlyGly-CTERM containing proteins by rhombosortase in Vibrio cholerae. PLoS Pathog 14:e1007341.
2. Shannon A, Johnson T, Roberts CS, Chaton CT, Korotkov KV, Sandkvist M. 2025. The PDZ domain of EpsC is required for extracellular secretion of VesB by the Type II secretion system in Vibrio cholerae. J Bacteriol doi:10.1128/jb.00144-25:e0014425.
3. Sikora AE, Zielke RA, Lawrence DA, Andrews PC, Sandkvist M. 2011. Proteomic analysis of the Vibrio cholerae type II secretome reveals new proteins, including three related serine proteases. J Biol Chem 286:16555-66.
4. Knauf VC, Nester EW. 1982. Wide host range cloning vectors: a cosmid clone bank of an Agrobacterium Ti plasmid. Plasmid 8:45-54.
5. Donnenberg MS, Kaper JB. 1991. Construction of an eae deletion mutant of enteropathogenic Escherichia coli by using a positive-selection suicide vector. Infect Immun 59:4310-7.
6. Sikora AE, Lybarger SR, Sandkvist M. 2007. Compromised outer membrane integrity in Vibrio cholerae Type II secretion mutants. J Bacteriol 189:8484-95.
7. Furste JP, Pansegrau W, Frank R, Blocker H, Scholz P, Bagdasarian M, Lanka E. 1986. Molecular cloning of the plasmid RP4 primase region in a multi-host-range tacP expression vector. Gene 48:119-31.
8. Waack U, Warnock M, Yee A, Huttinger Z, Smith S, Kumar A, Deroux A, Ginsburg D, Mobley HLT, Lawrence DA, Sandkvist M. 2018. CpaA Is a Glycan-Specific Adamalysin-like Protease Secreted by Acinetobacter baumannii That Inactivates Coagulation Factor XII. mBio 9.
9. Roberts CS, Shannon AB, Korotkov KV, Sandkvist M. 2024. Differential processing of VesB by two rhomboid proteases in Vibrio cholerae. mBio 15:e0127024.
10. Scott ME, Dossani ZY, Sandkvist M. 2001. Directed polar secretion of protease from single cells of Vibrio cholerae via the type II secretion pathway. Proc Natl Acad Sci U S A 98:13978-83.
